# History is written by the victors: the effect of the push of the past on the fossil record

**DOI:** 10.1101/194753

**Authors:** Graham E. Budd, Richard P. Mann

**Affiliations:** Dept of Earth Sciences, Palaeobiology, Uppsala University, Villavägen 16, Uppsala, Sweden, SE 752 36.; Department of Statistics, School of Mathematics, University of Leeds, Leeds LS2 9JT, United Kingdom.

**Keywords:** Crown groups, diversification rates, mass extinctions, the push of the past, molecular clocks

## Abstract

Phylogenies may be modelled using “birth-death” models for speciation and extinction, but even when a homogeneous rate of diversification is used, survivorship biases can generate remarkable rate heterogeneities through time. One such bias has been termed the “push of the past”, by which the length of time a clade has survived is conditioned on the rate of diversification that happened to pertain at its origin. This creates the illusion of a secular rate slow-down through time that is, rather, a reversion to the mean. Here we model the controls on the push of the past, and the effect it has on clade origination times, and show that it largely depends on underlying extinction rates. An extra effect increasing early rates of lineage generation is also seen in large clades. These biases are important but relatively neglected influences on many aspects of diversification patterns, such as diversification spikes after mass extinctions and at the origins of clades; they also influence rates of fossilisation, changes in rates of phenotypic evolution and even molecular clocks. These inevitable features of surviving and/or large clades should thus not be generalised to the diversification process as a whole without additional study of small and extinct clades.

## 1 Introduction

The patterns of diversity through time have been of continuous interest ever since they were broadly recognized in the 19th century (e.g. [1]). In particular, both major radiations (such as the origin of animals [2] or angiosperms [3]) and the great mass extinctions (e.g. the end-Permian [4] or end-Cretaceous [5] [6]) have attracted much attention, with an emphasis on trying to understand the causal mechanisms behind these very striking patterns. For example, the “Cambrian explosion” and “Great Ordovician Biodiversification Event” have both been discussed at great length, with mechanisms as diverse as the cooling of the Earth to bombardment with cosmic rays or secular changes in developmental mechanisms being suggested [7]. However, in the midst of this search, the effect of survival biases on creating the patterns under consideration have hardly been considered. The last decades have also seen a great deal of interest and work on mathematical approaches to diversification and extinction (e.g. [8] [9] [10] [11] [12]), including some that touch on the topics considered by this paper (for example, see especially [13] [14] [15] and [16]), but there is hardly any literature on the dynamics of clade origins from the perspective of survival biases and their effect on the fossil record. In this paper, then, we wish to explore the basis for such biases and then consider how it is exported to various important aspects of the observed large scale patterns of evolution, with particular (but not exclusive) focus on the sort of data that can be extracted from the fossil record. In the following analyses, we calculate over an interval of 0.1myrs and plot graphs with an interval of 2myrs. We term the (often unvarying) rate of a particular type of event (extinction or speciation) in a model as the “background” rate; and the rate of such events measured during a particular interval of time as the “observed” rate. For example, the background rate of rolling a six using an unbiased 6-sided die is 0.17; but if in seven trials five sixes happened to be rolled, the observed rate would be 0.71.

## 2 The “push of the past”

Nee and colleagues [17,18] summarised the general mathematics of stochastic birth-death models as applied to phylogenetic diversification (see also especially [15]). In such models, each lineage has a certain chance of either disappearing (“death”, the rate of which is usually labelled ‘*μ*’) ^or^ splitting into two (“birth”, rate ‘λ’). Many models that consider diversity in this way have used constant birth and death rates which have revealed much of interest about phylogenetic processes [19,20]. As Nee et al. pointed out [17], conditioning clades on survival to the present generates two biases in the rate of diversification through time: the “push of the past” (POTPa) and the “pull of the present” (POTPr).

The POTPa emerges as a feature of diversification by the fact that all modern clades (tautologically) survived until the present day. This singles them out from the total population of clades that could be generated from any particular pair of background birth and death rates: clades that happened *by chance* to start off with higher net rates of diversification have better-than-average chances of surviving until the present day. As Nee et al. ([21]) put it, such clades “get off to a flying start”. Once clades become established, they are less vulnerable to random changes in the observed diversification rate, and this value therefore tends to revert to the background rate through time. Long-lived clades thus tend to show high observed rates of diversification at their origin, which then decrease to their long-term average as the present is approached (c.f. [22]). A similar effect should apply to now *extinct* clades (such as trilobites) that nevertheless survived a substantial length of time. This effect is analogous to the “weak anthropic principle”, which contends that only a certain subset of possible universes, i.e. those with particular initial conditions, could generate universes in which humans could evolve in order to experience them. Similarly, to ask the question why living clades appear to originate with bursts of diversification that then moderate through time is to miss the point that this pattern is a *necessary* condition for (most) clades to survive until the present day.

The ‘pull of the present’ (POTPr) is, conversely, an effect seen in the number of *lineages* through time that will eventually give rise to living species, which is effectively what is being reconstructed with molecular phylogenies. As the present is approached, the number of lineages leading to recent diversity should increase faster than the background rate of diversification because less time is available for any particular lineage to go extinct. Thus, the POTPa affects reconstructions of *diversity* through time; and the POTPr affects the number of *ancestral lineages* through time (Fig. 1A).

Despite the theoretical considerations above, molecular phylogenies, far from showing a pronounced POTPr, typically show either no change in observed diversification rate as the present is approached, or even show a marked *slow-down* in rate – the opposite effect to what might be expected. Why this might be the case has been the subject of intensive research over the last few years, with various models being proposed, the most important of which are the ‘protracted speciation’ model of Etienne et al. 2012, and various proposals that carrying capacities of environments lead to ‘diversity-dependent diversification’ (DDD) (see [23] for review; see also [24] for skepticism about the reality of the effect). Diversification patterns thus potentially show two sorts of slow-down: one after the initial burst of diversification; and one as the present day is reached. This paper deals with the first of these effects. When clades had themselves a recent origin, these two effects can of course be confounded, so in our discussion we largely confine ourselves to old clades whose “beginning” and “end” are clearly separated.

Although the POTPa is known about (e.g. [25] [15]), emerges naturally from conditioning clade survival to the present in diversification simulations and has even been examined in some detail in a few cases [26] [22], it has had surprisingly little penetration into especially the palaeontological literature, where it is most important, and yet has been almost ignored. In this paper, therefore, we wish to show how influential an effect it and related effects are, and discuss its implications for general discussions about the reasons behind typical patterns of diversification seen in the fossil record and molecular phylogenies, including guidelines for how it and related effects might be detected.

## 3 Mathematical analysis

In this section we extend on the approach of Nee *et al.* by explicitly conditioning the birth-death model on the number of extant species in the crown group, and considering the full distribution of clade abundances over time rather than just the central expectation.

### 3.1 Estimating the number of extant and surviving clades through time

As noted by previous studies [17, 27], in the classic birth-death model with speciation rate per species *λ* and extinction rate per species *μ*, the number of extant species, *n_t_* in a surviving clade at a given time *t* from the origin at time zero, obeys the following zero-truncated geometric probability distribution

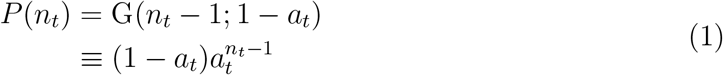

with

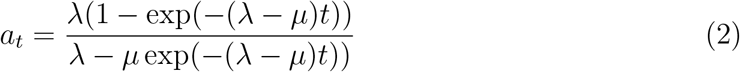

It is also useful to introduce the survival probability, s_Δ*t*_: the probability that a lineage with one originating species will survive for a duration of time Δ*t*,

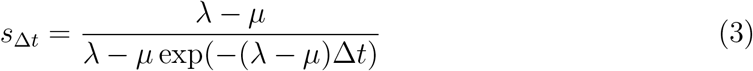

For the limiting case where *λ* – *μ* ⟶ 0, see [27]. Nee *et. al* proceeded by conditioning the distribution of *n_t_* on the tree surviving until some future time, *T*. Here we also condition on there being *n_T_* extant species at time *T*. By Bayes’ rule:

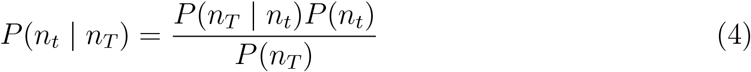

Two terms in this equation are given immediately from equation 1. We can evaluate the remaining term, *P*(*n_T_* | *n_t_*), by recognising *n_T_* as a sum of *m_t_* i.i.d. geometric random variables obeying equation 1 over a time period *T* – *t*, where *m_t_* is the (unknown) number of species at time *t* that will give rise to surviving lineages. This implies that *P*(*n_T_* | *m_t_*) follows a truncated negative binomial distribution (with *n_T_* taking a minimum value of *m_t_*):

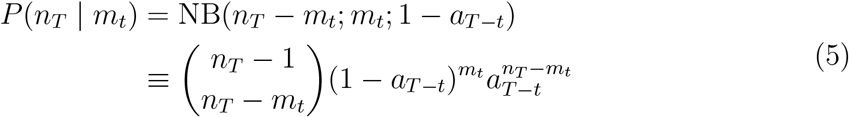

The number of lineages that survive depends on the probability, *s*_*T*–*t*_ for a lineage to survive from time *t* to time *T*, and follows a binomial distribution:

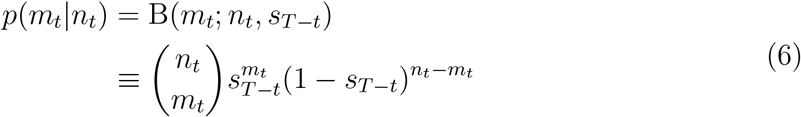

Combining the previous equations, and summing over the unknown value of *m_t_*, we are left with the following expression for the number of living species at time *t*, conditioned on the number of species in the present:

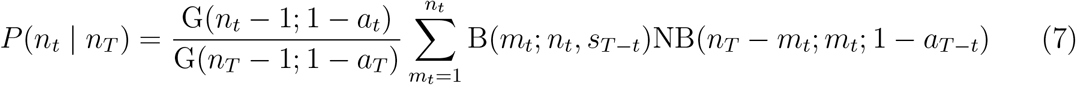

where the relevant probability mass functions are as defined above. We can further evaluate the conditional probability of *m_t_* – the number of species at time *t* which will have at least one descendant at time *T*. Here we integrate over possible values of *n_t_*:

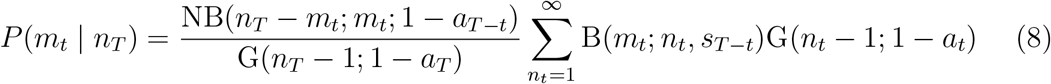

What sort of expectations should we have about the size and duration of the POTPa? By looking at a small initial interval of time Δ*t* from the origin and considering both the probability that the clade has diversified to two species in this interval and the probability that it will survive to the present, we can estimate the initial observed rate of diversification for surviving clades:

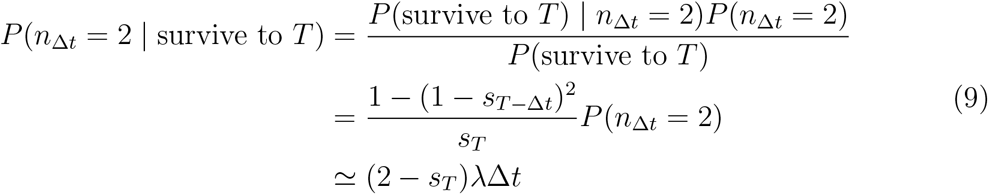

where we have assumed that *s*_*T*-Δ*t*_ ≃ *s_T_* for small Δ*t*. It follows that the initial rate of diversification, *R*_0_, in the POTPa can be estimated by:

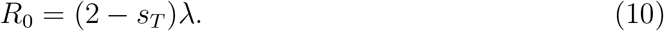

If we look back to the origins of major clades, we expect *s_T_* to be small for geologically significant periods of time, and thus for these examples the rate can be further approximated as,

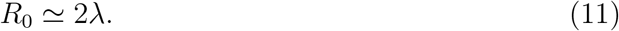

(we note that similar results concerning the interior branch lengths of reconstructed trees have been derived by [14]).

It is important to note that at the precise origin of a clade that will survive to the present, the observed *extinction* rate is necessarily zero, since any extinction event would terminate the clade. Thus at this point the observed speciation and diversification rates are the same.

What sort of expectations should we have about the size of the POTPa? The result above shows us that if we know *λ* and *μ*, we can immediately calculate that the expected POTPa should produce a decline in observed diversification rate from about 2*λ* as an initial value down to *λ* – *μ*. as deduced from the fossil record. However, the broad confidence intervals on this value place the 95% range on this value widely: for example, in Fig. 1 over the first million years, the initial rate could be as low as 0 and as high as 3 (ie 6*λ*). Thus slowdowns seen in rates of diversification that begin with a wide range of values and quickly decline (largely being over by the time of the establishment of the crown group), with reconstructed rates in the stem lineage being significantly higher than in the generated plesions, are attributable to the POTPa. For the case of the birds given above, one would expect (in the fossil record) an observed initial diversification rate of about 1.25, ie about 20 times faster than the background rate. Another example is provided by the study of diversification rates in placental mammals [28] which simulates a best fit homogeneous model with parameters *λ* = 0.7, *μ* = 0.6 and *λ* – *mu* = 0.1. A POTPa effect would generate a decline in rates from 1.4 to 0.1 over 5-10 million years, which closely matches their reconstruction of phenotypic rates (that correlate in their data to diversification rates. Thus, the calculated POTPa corresponds closely to real-world examples.

Even though the POTPa as an *average* effect quickly declines in time, it must persist along the surviving lineages: every clade that will survive to the present commences with one original species, which is vulnerable to extinction. As the survival rate typically remains low until close to the present, it follows that the constant renewal of the surviving stem lineages follows a quasi-fractal pattern of repetition. A further notable feature is that as the present day is approached, and the survival probability thus tends to one, the observed rate of speciation along surviving lineages declines back towards λ.

An example plot displaying the various parameters that govern this analysis is given in Fig. 1A. (c.f. [17]). This plot is for a clade that has 10,000 living species/lineages; which emerged about 500 million years ago, and which has an average lineage duration of 2 million years (i.e. *μ* = 0.5).

The blue line gives the number of species at any particular time; and the slope of the blue line is the observed diversification rate governed by the time elapsed and total number of taxa at the Recent, *n_T_*. Conversely, the red line gives the number of lineages that gave rise to living species/lineages. We take as the rate of speciation the maximum likelihood estimate of *λ*, given *μ,T* and *n_T_*, which in this case is 0.5107. Thus the rate of diversification (*λ* – *μ*) is c. 0.0107 per species per million years. As can be seen, the slopes of the two lines diverge at the beginning, representing the push of the past, and at the end, representing the pull of the present, both of which are large in this case. If there had been a *deterministic* (i.e. non-stochastic) radiation of species from the Cambrian onwards with the net diversification rate of 0.0107 species per species per million years, then instead of 10,000 there would have been only about 210 species of this taxon today. Fig 1B shows the implied large spike in the initial observed diversification rate, owing to the POTPa, with the initial observed diversification rate (~ 1) being 100 times the underlying average. Note also that this effect generates a (non-causal) correlation between diversity and diversification rate (Fig. 1C): as diversity increases, (average) rate of diversification decreases.

How realistic are the numbers in our example? The size (and thus importance) of the POTPa depends on the rate of extinction relative to the other parameters. Extinction rates have proven difficult to estimate from both molecular phylogenies (notably [29]; but see also [30] and [31] and the fossil record (see e.g. discussions in [32] [33] [34] [35]). Nevertheless, the fossil record in particular shows that extinction rates must be relatively high, as most species across a wide range of taxa only last a few million years at most in the record (e.g. [36]). For example: extinction rates over all marine invertebrates have been broadly estimated at c. 0.25 per species per million years [37] [38], and for Caenozoic mammals at up to 2 per species per millon years [37, 39]. Even highly conservative estimates of background extinction rates, which partly equate species with genera in the fossil record, suggest rates in excess of 0.13 ([40]; cf. [41]; see [32], however, for a critique of the methodology used in the latter, which produces a notable downwards bias). An example clade would be the birds, that, although may have had their crown group origin some 120 Ma ([42]; but see also [43] and [44] for a more compressed view of bird evolution), nevertheless seems to have undergone a mass extinction along with the other dinosaurs 65Ma [45]. Such a clade thus took approximately 65-70Myrs to radiate into 10,000 species. Assuming a (probably conservative) extinction rate of 0.5 based on other land vertebrates [46]; this would imply a speciation rate of approximately 0.62 and a diversification rate of c. 0.12 (c.f. [42]). Of course, all these numbers are approximate, but our aim with them is to show that the patterns we discuss in this paper arise from very typical empirical values seen in analyses of extinction and diversification. Assuming that birds are a “typically” sized clade (see below), this would imply a notably enhanced rate of diversification with initial rate of c. 1.2 species per species per million years that would decline over about 5-10 million years (c.f. [44]).

Although the parameters we have explored in this paper thus seem to be typical of diversifications over a large range of species numbers and time, we wish to stress the important point that the clades that emerge from them that survive for long periods are rare. In our example, although the living clade has had to survive for 500 Myrs, the median survival time of an average clade generated by these parameters is only 2 Myrs. Furthermore, only 2.1% of clades thus generated will survive for 500Myrs. These numbers emphasise how unusual long surviving clades are, even when there is a net positive diversification rate: survival rates for other diversification scenarios are given in Fig. 4. The POTPa generally has the paradoxical effect of making high extinction rates *increase* observed rates of diversification and numbers of living species - in the rare clades that managed to survive.

**Figure 1:**
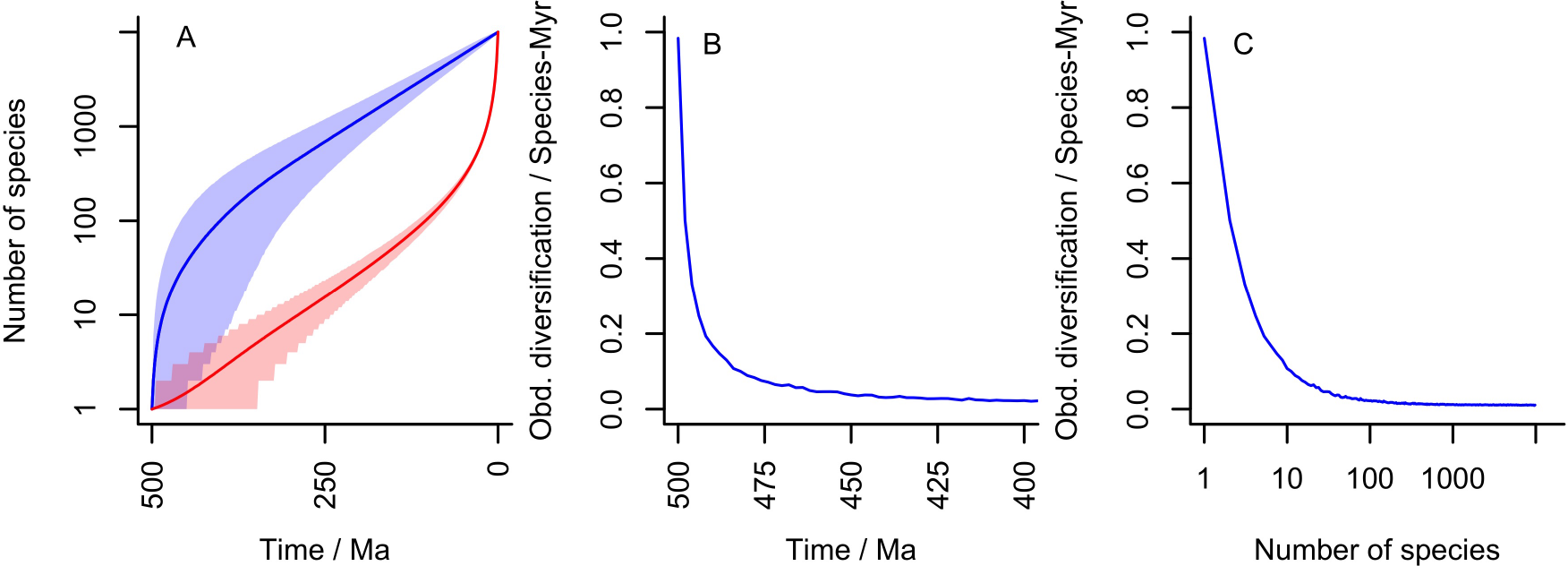
An example diversification with 10,000 living species, an extinction rate of 0.5 per species per million years and a diversification time of 500 Myrs. The implied speciation rate is 0.5107 per species per million years and thus the underlying diversification rate is 0.0107 per species per million years. **A** Diversity plot through time. As in all other figures, the blue line is the number of species at time *t* and the red line the number of species that will give rise to living species. Shading gives 95% confidence areas. Note large POTPa and POTPr. **B** Observed diversification rate at beginning of diversification (note scale of 100 Myrs). **C** Implied diversification rate correlation with diversity generated by this distribution

### 3.2 Crown group origins

If the number of species at time *T* is known (*n_T_*), and if the number of lineages that will survive at time *t* is also known (*m_t_*), we can calculate the probability, *W*(*t*), that two randomly chosen species in the Recent will have a common ancestor at time *t*. This is the definition of a “randomly selected” crown group used by Raup [47]. We first need to pick the first species at random (with probability one), then we need to pick a second species that has the same ancestor at time *t* - this must first be one of the *n_T_* – *m_t_* remaining species that do not inevitably have to join up with the other *m_t_* – 1 ancestor species, and then a randomly selected species from this remaining set will have a 1 */mt* probability of sharing an ancestor with the first selected, thus:

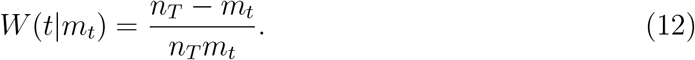

Accounting for the uncertainty in our knowledge of *m_t_*, our estimate of *W*(*t*) requires a posterior-weighted summation over the possible values of *m_t_*:

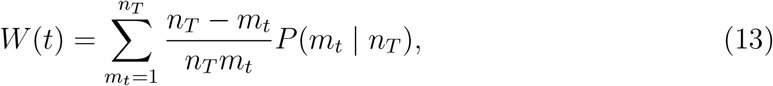

where the posterior distribution on *m_t_* is calculated as above. *W*(*t*) represents a cumulative distribution function for the timing of crown group origins for randomly selected pairs of species, looking backwards in time. The corresponding probability density function, *w*(*t*), is given by differentiation of *W*(*t*):

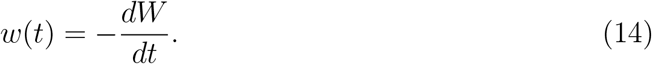

Compared to Raup’s model (which can be most closely approximated by the Yule process, although he did not include a stochastic component) our model delays the average time of origin of Raupian crown groups because of the effect of the POTPr of allowing a longer period of lower early lineage diversification rates. Nevertheless, it remains true that randomly-selected pairs of taxa will also tend to have early origins (Fig. 4F, I). As can be seen (Fig. 4C), the Yule process forces crown-groups defined in this way to emerge very early. Budd & Jackson [48] simulated the origin of the *first* crown groups in clades conditioned on survival (c.f. [22]), a topic of much interest in “Cambrian Explosion” literature (e.g. [49]).In simulations that start with one lineage and go on to diversify to the Recent, the time the simulation begins can be taken as the origin of the total group, and the emergence of the crown group (for the entire clade) when *m,_t_* (the number of lineages at any time *t* that will give rise to living descendants) is equal to two (i.e. the basal split of the crown group is formed). Since this state can only be reached from a previous state of m_t_ = 1, the probability density *u*(*t*) that the first crown emerges at time *t* can therefore by calculated by considering the rate of change in the probability that *m_t_*=1:

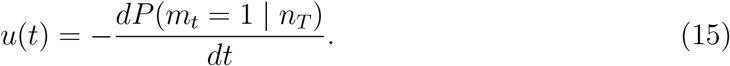

As summarised in [17] and Fig. 1A, we can see that *m_t_* in the early stages of diversification essentially depends on *λ* – *μ*: *m*_*t*_ ≃ exp([*λ* – *μ*]*t*). Thus a simple approximation for the expected length of time it takes for the first crown-group to emerge is given by:

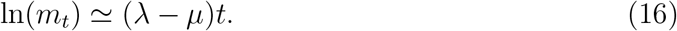

Thus *t_cg_*, the time in millions of years ago that the first crown group is expected to have emerged is simply

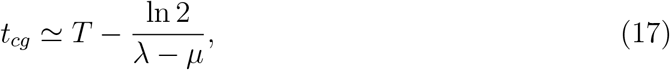

where *T* is the time elapsed since the origin of the total group. As an interesting aside, the underlying diversification rate *λ* – *μ* is thus approximated by:

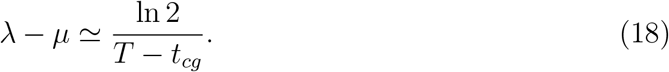

The combination of the POTPa and the dependence of *t_cg_* on *λ* – *μ*, means that stem and crown groups exhibit different characteristics of diversification and diversity, as the first crown group tends to emerge as the effect of the POTPa fades away. An example of this is given in Fig. 2B.

**Figure 2:**
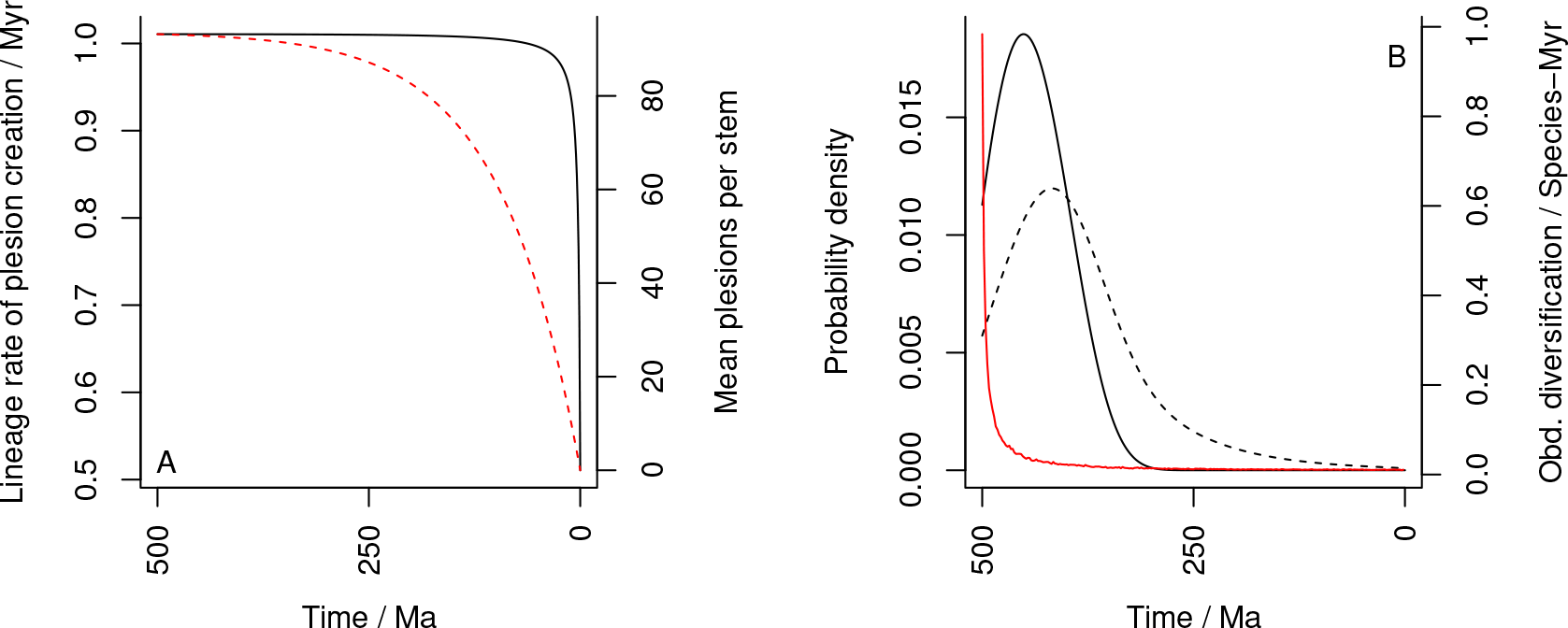
**A** Illustration of the rate of plesion creation along the surviving lineages (black solid line) and the mean number of plesions created along each stem group (red dashed line) through time. As rate of plesion creation is almost flat for most of the time, it follows that the decline in number of plesions per stem group depends on the stem groups decreasing in size temporally. **B** Observed diversification rate (red) and probability density functions of the first crown group (black, solid) and origin times for pairs of random living species (black, dashed) against time. All plots for a diversification over 500Myrs in total, *n_T_* = 10,000, *λ* = 0.51 and *μ* = 0.5 (i.e. the same example as Fig. 1). Note the likely emergence of the first crown group as the POTPa decays.

## 4 The push of the past and the fossil record

### 4.1 Overview

Within a particular total group, then, stem groups are characterised by high observed diversification rates and low diversity; and crown groups by low diversification rates and increasing diversity (c.f. Fig. 1C). The interaction between the crown group and the POTPa allows us to understand why it is that the crown group emerges just as the POTPa dies away: the POTPa is an effect seen when there are few (surviving) lineages and as soon as there are two rather than one, the likelihood of the clade surviving until the present is considerably increased.

It is important to note that the high rates of observed diversification in stem groups are not *general* features, as we are applying a homogeneous model of diversification. Rather, unusually high observed diversification rates are concentrated in the stem *lineage* that leads to the crown group(s) (c.f. [14]). Stem groups should thus generate a high number of so-called “plesions”(i.e. extinct sister groups to crown groups [50] [51]) which themselves will diversify and go extinct at approximately the background rate governed by *λ* – *μ*. From equation 10 we can see that the rate of speciation along most of the stem lineages, and thus the rate of production of plesions, remains close to 2*λ*, although the rate slowly declines until close to the Recent, when it precipitously drops to *λ*. Similarly, lengths of stem-groups also decrease, over a longer timescale, as the present is reached (see Fig. 2A for graphical treatment).

**Figure 3:**
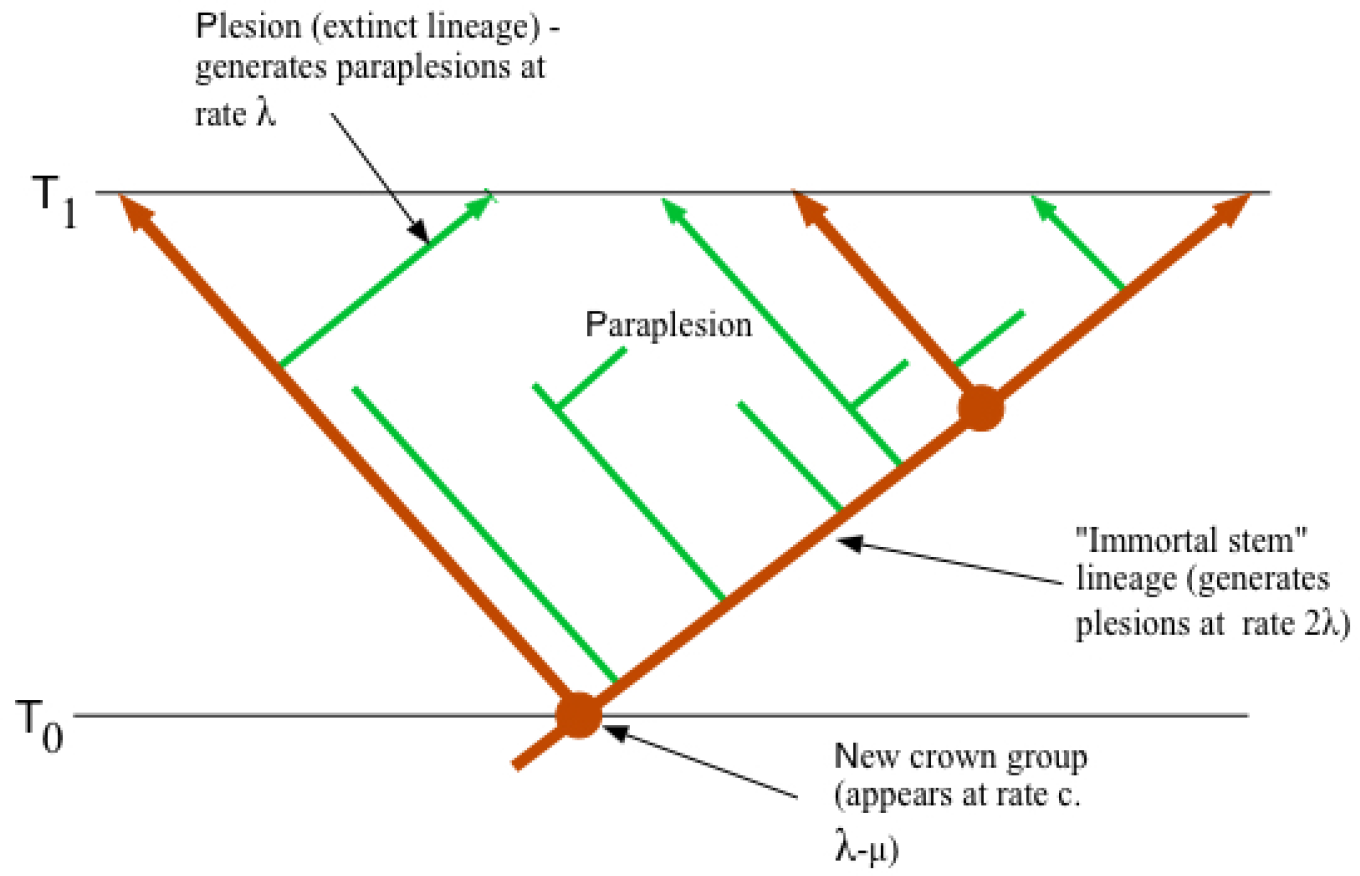
A small section of a tree at a time distant from the present. Red branches represent lineages that survived until the Present, and where they diverge represents the birth of a new crown group. Green branches represent plesions that do not survive until the present. As per the results presented herein and in [14], the stem lineages species generate plesions at a rate close to 2*λ* and the plesions themselves speciate at rate *λ*: the crown groups form at rate *λ* – *μ*.

This analysis gives us a remarkable perspective on the fossil record (Fig. 3), which is after all considered on methodological grounds, as taxa in cladograms are only ever terminals, to be composed entirely of plesions (Budd 2003). Average rates of speciation (and, as we shall argue below, rates of phenotypic evolution) typify the clouds of plesions that are constantly being generated (and dissipating) at a high rate; but underlying them, and hidden from view, are stem lineages that speciate at twice the normal rate. It is only briefly, at the beginnings of radiations and after the great mass extinctions (see below) that these obscuring clouds are stripped away, and we get to peer at the underlying hyperactive stem lineages. Once again though it must be stressed that this pattern only emerges as a result of our perspective in the Recent, which allows us to distinguish stem lineages from plesions.

### 4.2 Diversification scenarios

Armed with the mathematical analysis and example above, we are now in a position to analyse various scenarios that might play out in patterns of diversity and the fossil record. In each, we wish to examine: i) the size of the POTPa effect; ii) the distribution of the timing of crown group origins and iii) the relative proportions that the stem and crown groups take up of the total group.

### 4.3 The Yule process

The Yule process [52] governs diversification processes with no extinction, i.e. that *μ* = 0. Of course this is not realistic over geologically-significant time periods, but nevertheless is important to show the contrast between this and more realistic models. Furthermore, it can be used to model surviving lineages through time, that have no extinction.

Under the no-extinction model, as all species give rise to living lineages, it is clear that the blue and red lines of Fig. 1 are coincident (Fig. 4A) irrespective of the error on each. There is neither a pull of the present nor a push of the past (Fig. 4B), and the slope of the line simply gives the diversification rate through the time required to lead to the observed *n_T_*. The rate of diversification is completely constant along the mean, since the diversification rate has been selected to generate *n_T_* (ie 1000 species in this case). Nonetheless, as the confidence region shows, early fluctuations in this process are possible, which we consider further later. Another feature of the no-extinction model is that total and crown groups are nearly coincident for any particular clade, as stem-groups grow by extinction [53]. A lag at the beginning is possible though, before the first speciation event takes place.

### 4.4 Models with net diversification and extinction

Fig. 4D-F and G-I model two net diversification models; one with *μ*, = 0.1 and the other with *μ* = 0.5, both with the best-fit implied *λ* (the maximum-likelihood value given *T*, *n_T_* and the selected value of *μ*). As can be seen, increasing *μ*, increases the POTPa.

If *μ* is set *very* low (e.g. *μ* = 0.01 for *T* = 500 Myrs and *n_T_* = 1000), then the POTPa can be much reduced. However, such models imply very implausible species longevities (a typical species would be expected to survive 100 Myrs in this model, numbers that are not realistic for the Phanerozoic (they may, however, be more appropriate to the Proterozoic ([54]).

**Figure 4:**
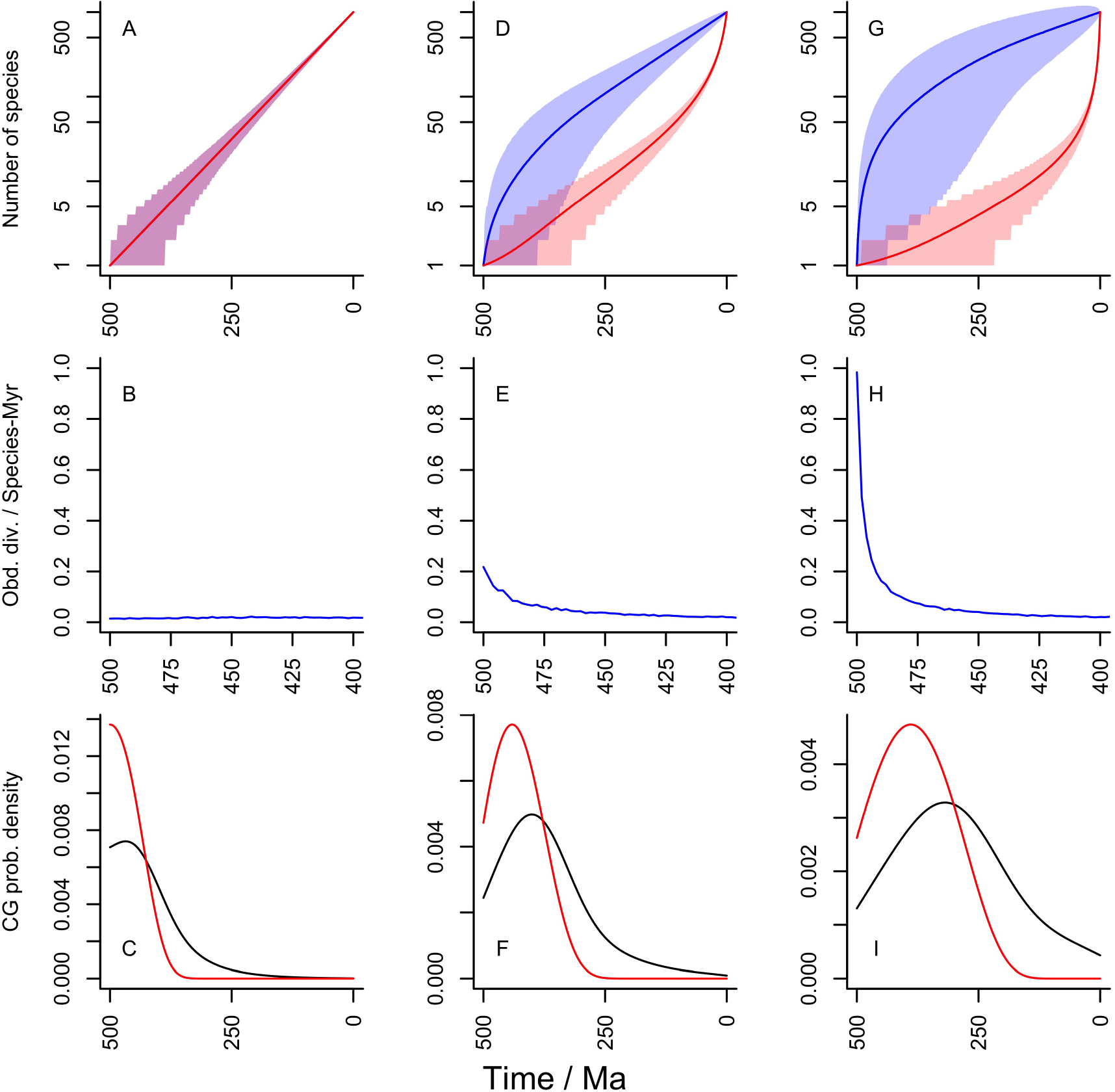
Patterns of diversification and stem- and crown-group formation for different (constant) diversification parameters. For each column T= 500Myrs and *n_T_* = 1000. All rates given per species per million years. Shading gives 95% confidence areas. Row one gives plots of diversity and diversity that gives rise to extant species through time; row two gives observed average diversification rates through time over the first 100Myrs (ie. the POTPa effect); row three gives probability density function plots for the appearance of the first crown group (red) and for crown groups defined by random pairs of living species. **A-C:** Yule process with *μ* = 0 and *λ* = 0.014. **D-F:** low *μ* net diversification with *μ* = 0.1 and *λ* = 0.109. **G-I:** high *μ* net diversification with *μ* = 0.5 and *λ* = 0.505. For the second column, median clade survival time is 10.5 Myrs and 8.2% of clades would survive 500 Myrs; for column 3, the corresponding numbers are 2 Myrs and 1%. Note different time scale on second row.

Models with the same background diversification rate can have high and low turnover (e.g. *λ* = 0.6, *μ* = 0.5, or *λ* = 0.2, *μ* = 0.1). As models with larger diversity fluctuations in will be more vulnerable to extinction than ones with small ones, it follows that high turnover scenarios require a larger POTPa in order to escape the early period of vulnerability.

## 5 Mass extinctions

We have chosen for simplicity a diversification model that is diversity-independent and has homogeneous rates of extinction both through time and for taxon-age (c.f. [55]). Nevertheless, the handful of mass extinctions through time have had a large impact on diversification patterns [56]. The most important are perhaps the end-Permian (with c. 80% of all species going extinct [57] and the end-Cretaceous (c. 68% loss [57]). Such events could be considered as simply “resetting the clock” - i.e., if evidence exists that extinction was extremely severe in a particular clade, then T should be considered to restart at that point. Some overall patterns of diversification suggest that the only truly important mass extinction in this regard is the end-Permian one [58] [59], which divides Phanerozoic time more or less into two, with large POTPa, but largely uncommented, effects at the beginning of each. One interesting effect is that the bigger a mass extinction, the bigger the subsequent POTPa would be, assuming *something* survives to the present. Even so, these big pushes can never make up for the lost diversity, even if they compensate for it to some extent. For example, for a diversification that started 500Ma, and that would have generated 1000 living species without any disturbance, and with a background extinction rate of 0.5 (and implied maximum-likelihood speciation rate of 0.504), a mass extinction 250Ma down to only one species and the subsequent POTPa and re-radiation would only generate 240 living species - it is a rerun of the original radiation but in half the time. On the other hand, without any POTPa, this re-radiation would be expected to generate only 3 living species.

It is possible to model the POTPa with a standing diversity, and show how the size of the POTPa declines as surviving diversity increases. We modelled this by plotting number of survivors against immediate observed diversification rate post-extinction (Fig. 5) for different rates of background extinction for a radiation that took 250Myrs to generate 1000 species. As can be seen, extinctions can indeed generate large POTPa, but the number of remaining species for the clade needs to be reduced to a few percent of their original numbers. Thus, large POTPa effects after a mass extinctions are likely to be contingent on large extinctions preceding them in the clade in question.

Another POTPa-like bias may also be having an effect, which is the effect of the POTPa on fossilization rates themselves. One of the controls on the preservation probability of a taxon is its true (as opposed to fossil record) temporal duration, and thus its extinction rate ([60]; [61]). When diversity drops to a low level, survivorship over the next short interval of time is compromised, with the implication that only taxa that experience unusually high rates of diversification are likely to survive - and thus enter the fossil record. 6 shows there is a strong relationship between survivorship on a million or sub-million year scale and diversification rates. In brief: taxa straight after a mass extinction or at the beginning of a radiation have an unusually poor chance of entering the fossil record, as their diversity is so low and their chance of almost instant extinction is so high. However, the taxa that by chance experience high rates of early diversification are much more likely to survive long enough to generate a discoverable fossil record. Such an effect may at least partly lie behind the observation that fossilization rates seem to be depressed after mass extinctions (notably the end-Permian [62]). Thus, one interesting aspect to this pattern in the record, that such “recoveries” seem to be delayed, with clades sometimes taking millions of years to show increased rates of diversification (see e.g discussion in [58]), may be partly explicable by this effect too: early survivors are simply such low diversity that they tend to go extinct faster than they can enter the fossil record.

**Figure 5:**
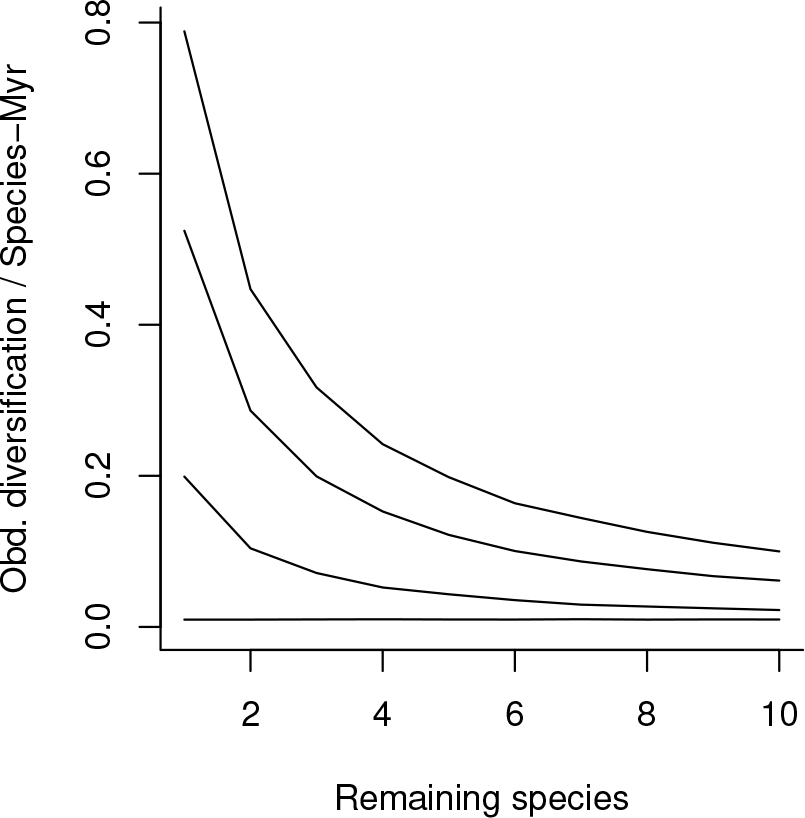
The impact of number of remaining species on a post-extinction POTPa on a clade that had diversified for 250Myrs to generate 1000 species, assuming clade survival to the present, with baseline diversification *λ* – *μ* = 0.01. The curves from bottom to top represent background extinction rates of *μ* = 0 (Yule process), 0.1, 0.3, and 0.5 per species per million years.

### 5.1 The “Copernican” nature of Birth-Death models

The various cases we have considered above show that the POTPa is in general a very important factor that cannot be neglected in trying to understand diversity patterns of the past. The most important control on the size of the POTPa is the extinction rate (compare Figs 4E and H) although time to the Recent also has some effect. Thus, when significant time periods have passed, the POTPa is always large unless the background extinction rate is extremely low (cf. [15])- much lower than seems to be typical for at least Phanerozoic taxa, which typically have a life time of a few million years ([58]).

Because of the nature of the homogeneous model we are using, we wish to stress its ‘Copernican’ aspect, i.e. that diversification is on average the same at all times. Each stem lineage will be characterised by a high POTPa, but as diversification continues, its distorting effect on *average* diversification rates in surviving lineages will be diminished by two factors. The first (which is small until the POTPr is reached) is that as time advances, each lineage has less time to survive until the present. The second is that as diversification proceeds, more average or even below average diversification-rate lineages will be present, and thus the overall average rate of diversification will be swamped by their diversification rates. In the first stem group, so few lineages are present that the implied POTPa on the stem lineage will have a disproportionate effect on average diversification rates. Such controls produce the characteristic decline in average actual diversification rates through time, even though an observer at any particular time would not notice any difference whatsoever.

**Figure 6:**
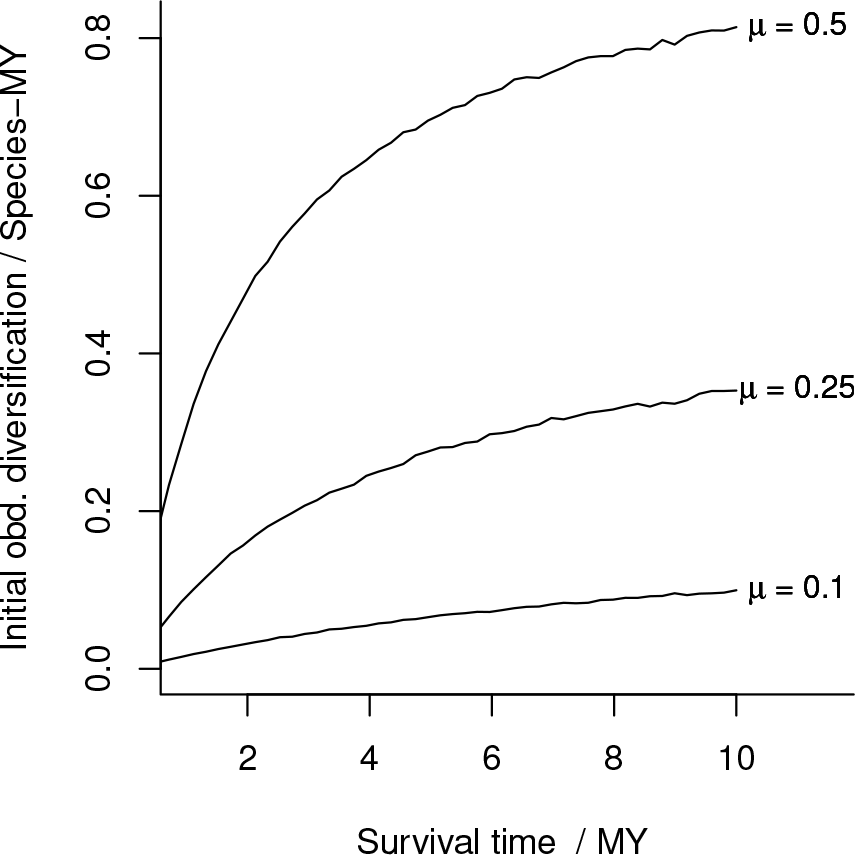
Expected observed initial diversification rate as a function of clade survival time for different values of underlying extinction rate, *μ* in a neutral model. As *μ* increases, a bigger and bigger POTPa is required to ensure the clade survives the first few million years.

## 6 The large clade effect: an analogy to the POTPa in reconstructed phylogenies

So far, we have considered the effect of survivorship biases in the blue line of Fig. 1. Our exploration of the POTPa shows, however, that when conditioned on survival, it remains nearly constant at along the *surviving* lineages at close to 2*λ* until the Recent is approached. Hence, it largely cannot account for long-term declines in phenotypic and molecular rates along the lineages (see below) or lineage production rates themselves. However, survival alone is not the only characteristic of a clade that can lead to statistical biases. As we have shown, the birth-death model can also be conditioned on the number of extant species in the Recent, *n_T_*. Therefore we can ask about the characteristics of outliers within the set of surviving clades, specifically those with a larger-than-expected present diversity [15]. Such outliers represent those clades which are held up as the most ‘successful’ examples of their type and, erroneously as we shall see, are often presented as ‘representative’ of their particular time of origination. As [63] observed through simulation analysis, larger than average clades are statistically more likely to show a slowdown relative to smaller clades. To illustrate this, we recalculated the example shown in Fig. 1, but conditioned it to generate 100,000 instead of 10,000 species (Fig. 7). Under such rare conditions, more lineages need to be generated than normal, and the most likely moment to do this (as can be seen in the confidence regions of Fig. 1A) is at the beginning, when overall numbers of lineages are small and statistical fluctuations more noticeable in effect.

**Figure 7:**
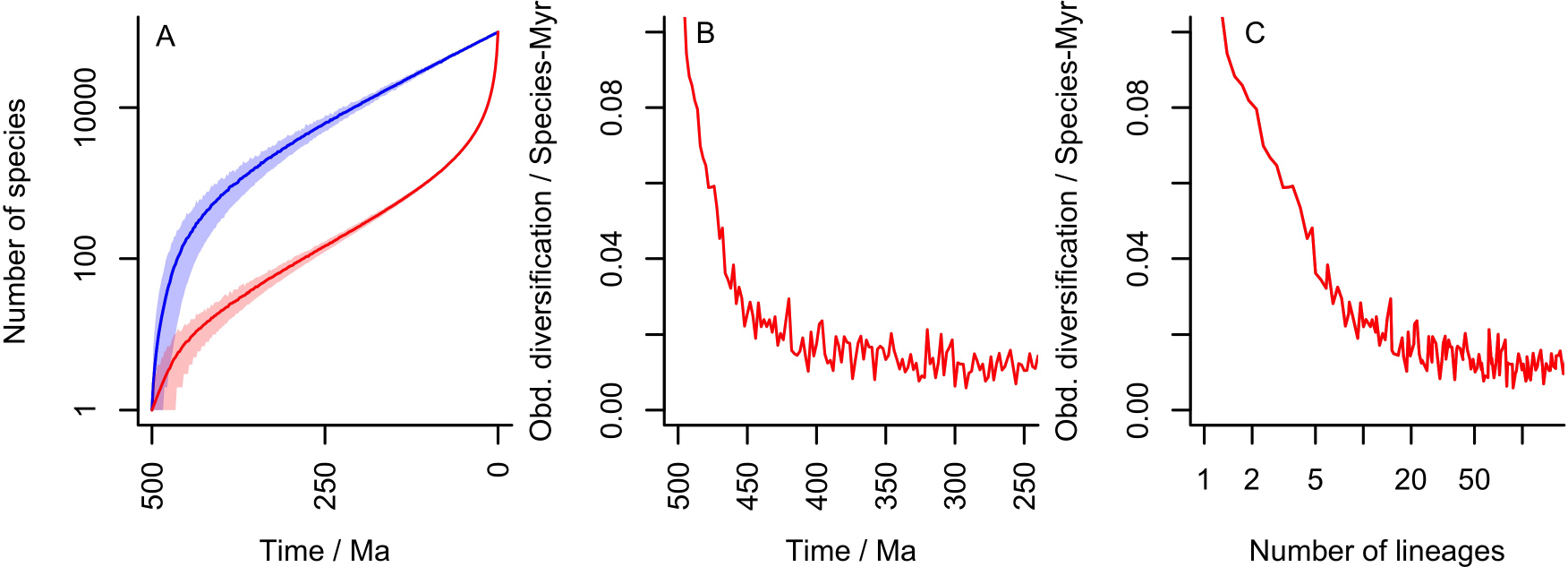
**A** Diversification when an exceptionally large clade is generated with the parameters of Fig. 1 (here, 10x larger than expected). An early lineage effect is introduced as this is where fluctuation in actual rates is most likely. **B** Calculated decline of lineage rate (thus in red) through time. Note that it lasts considerably longer than the POTPa although is of smaller effect (here initially c. 10x the background rate of *λ* – *μ*). The variations are owing to Monte Carlo numerical integration of equation 8, because of low overlap between the prior distribution and likelihood function for *m_t_*. **C** Implied lineage diversification rate correlation with lineage diversity generated by this distribution

Under such circumstances, a *lineage* effect is produced [22, 63], which could be called the large clade effect (LCE). Although it is smaller than the classical POTPa (in the example of Fig. 7 the rate of speciation along the lineages increases to 2.2λ from 2λ), it has the effect of speeding up the appearance of new living lineages near the beginning, and thus makes crown groups emerge (even) earlier. The lineage through time plot thus takes on a characteristic inverted “S” shape that is often seen in plots of molecular evolution (e.g. [12] [64]). Like the POTPa, it has a quasi-fractal organisation – within a given clade, larger sub-clades will experience greater early diversification than smaller sub-clades. As we discuss below, this effect thus influences rates of evolution in large clades, and will be particularly prominent if such clades have happened to attract more than average attention, as has indeed been suggested [15]. A correlation with lineage diversity is also generated Fig. 7C, which in our example can be seen for about the first 20 lineages.

The initial magnitude of the LCE can be explicitly calculated in terms of the relative magnitude of the clade relative to its expected size conditioned on the background speciation and extinction rates, E (*n_T_*| survive). To determine the initial rate relative to the background value *R* = *λ* – *μ*, we consider the probability that the new clade with one lineage diverges into two lineages within a small unit of time, both *a priori* and conditioned on the final clade size. Recalling that the distribution of *n_T_* conditioned on *m_t_* is negative binomial, we have:

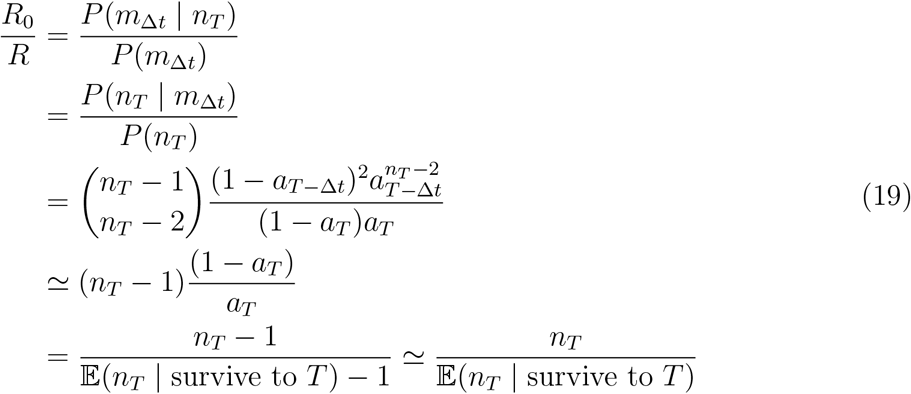

Thus we can see that the expected size of the initial LCE, and thus the magnitude of the later slowdown, is proportional to the eventual clade size. It should be noted the clade containing any randomly chosen species is expected to be twice the average clade size, and thus there is a consistent bias towards this effect appearing.

## 7 Effects of the POTPa and LCE on rates of phenotypic and molecular change

The rate of phenotypic change through time is another pattern that has seen a great deal of interest (e.g. [65]; [66]; [67] [68] [69] [70]; but see also [71]). A classical pattern of rates of phenotypic change is that rates are elevated at the origin of a clade and then show an exponential decline (e.g. [49]). Such a pattern looks, of course, like a POTPa effect, but this effect would seem to rely on a correlation between rates of phenotypic change and diversification. Whilst this seems both intuitively reasonable and has much theoretical backing, this pattern has been difficult to demonstrate and indeed some studies have failed to reveal it (e.g. [72]; [73]; but see also [74] who review the topic in general). Our model can account for such patterns by considering the fossil record to consist of plesions that are generated by a rapid rate of speciation in an underlying but unseen stem lineage. In principal at least, each of these speciation events (at least as recognised in the fossil record) should be accompanied by a set of diagnostic synapomorphies that accumulate within the stem lineage twice as fast as they do in the plesions that arise from it.

As the POTPa remains more or less constant along long stretches of the lineages until the present is reached, it follows that it should not generate a “slow-down” in measured rates of either phenotypic or molecular change along the lineages (ie measured along the red line of Fig. 1) when the present is far away. Average phenotypic rates of change should however decline through time when measured over all fossil taxa (ie measuring the rate of phenotypic change in the blue, rather than red, line of Fig. 1). The notable study of lungfish evolution through time by Lloyd *et al.* [75] reconstructed rates of phenotypic evolution through time and indeed noted such a decline (see their fig. 4; note that it also shows characteristic post-extinction spikes). However, as the authors note, the (reconstructed) stem lineage leading up to the extant lungfish retains high rates of phenotypic change much later than the initial rapid decline in overall rates, whilst the plesions appear to show no such pattern (the authors do not differentiate between the two in their analysis of the decline in rates). This pattern is exactly what the model we develop here would predict, as it confines the POTPa to the stem lineages, and suggests that their documented decline of phenotypic rates of evolution is a striking consequence of the POTPa (compare their fig. 1 with our Fig. 3). Clearly, it would be possible to test this pattern in other groups too. The POTPa should not, however, affect rates of molecular evolution because this cannot be measured in the blue line, only in the red.

The LCE, conversely, should affect rates measured along the lineages, but in a subtle way. For a big LCE, e.g. the 10x larger-than expected effect of our Fig 7, the rate of speciation along the lineages is initially increased only very modestly, i.e. in this instance from 2*λ* to 2.2*λ*. However, the initial rate of appearance of lineages increases from *λ* – *μ* to (*λ* – *μ*). The implication of both together is that although *rates* of speciation (and, by extension) molecular and phenotypic change hardly increase as a result of the LCE, the amount of that change in large clades that is *curated* into the present along the lineages is disproportionately sourced from the early stage of the clade’s history, when the LCE is in effect, at a rate proportional to the size of the LCE. Thus, the initial change per unit time per lineage should be increased proportionally to the size of of the LCE for both molecular and phenotypic change. This can give rise to large effects, which can be seen in the study of Lee et al. [67], where large initial rates for both phenotypic and molecular evolution can be seen as measured along the lineages. We note that the initial effect seen in Lee et al. is approximately 10x the normal rate and lasts for about 17 lineages, very similar to our calculation in Fig. 7C.

One implication of this finding is that in unusually large clades, one should expect a concentration of rapid molecular evolution in early lineages, and, if not corrected for, will create the effect of making molecular clocks overestimate origination times. Such an effect could in principle account for the continuing discrepancy between molecular clock estimates for the origin of the animals and the fossil record, for example [67] (but only in large clades, such as the arthropods [67] - and, of course, the animals themselves). Thus, although various studies have shown that rates of molecular change are at least loosely correlated with diversification rates (e.g. [76]; [77]; [78]; [79]; [80]), our model suggests that it is not diversification rates per se, but rates of lineage creation, that is correlated with (curated) amounts of molecular change.

## 8 Rate heterogeneity

A question that arises from this analysis is: ‘when is it appropriate to attribute heterogeneities in frequencies of events to intrinsic survivorship biases such as the POTPa and LCE rather than to adopt models where background probabilities vary exogenously through time?’ To take an example from the fossil record: some extinct taxa, notably the trilobites (e.g. [81]), indeed show a very rapid initial diversification, followed by a fairly drawn-out decline and final extinction. It is clear that such a decline cannot be realistically modelled by keeping the same background rate of diversification through time - it implies that the most appropriate background diversification rate has actually turned negative (c.f. [82]). One should note here however, that given that the trilobites experienced several mass extinctions, these singular events may have successively reduced their diversity to the point where they did become vulnerable to stochastic extinction, even with net positive diversification rates.

Several recent software packages (e.g. BAMM [83, 84] and MEDUSA [85, 86]) have been developed for detecting statistically significant rate shifts of this sort ([16]; see also [87] [42] for examples of their employment in different clades). How do the effects we outline here intersect with them? We have shown above the expected sizes of both the POTPa and LCE, which are themselves rate heterogeneities that arise from homogeneous models, when conditioned on either/or survival and clade size. Nevertheless, there is a difference between such heterogeneities and those seen from more complex models, because the heterogeneities that emerge from survivorship bias are strictly local as opposed to global in effect. The POTPa should leave a clear signal in the fossil record, because the expectation is that its rate heterogeneity should be confined to long-lived lineages: short-lived plesions should show no such pattern. Furthermore, such heterogeneity would be expected to be long-lasting and thus not be time-variant along the lineages until close to the present. Conversely, global rate heterogeneity, including forms that are diversity dependent, should affect all clades including short-lived ones, and this effect will be strongly time-variant. Rate heterogeneity that actually depends on (for example) diversity would be expected to affect all lineages including extinct ones. The signal of the POTPa should thus be readily detectable in suitable fossil data sets, i.e. ones from clades with a phylogenetic framework (e.g. [66]). With only raw diversity data through time of a particular surviving clade, any inferred rate variation from it that falls within the expectations of the POTPa outlined here should not be generalised as pointing to a time-specific period of enhanced diversification, e.g. after mass extinctions. Furthermore, given such patterns are inevitable, they should not be taken on their own as evidence for a particular generative mechanism.

The LCE, conversely, can by definition only be measured in the lineages. We have given an expression above for its expected size, which depends on the size of a clade relative to a base-line expectation for a given background rate of diversification. We note here, however, that there are two problems with estimating clade size. The first of these is relatively straight-forward, and relates to our inability to count all living species. This has been accounted for by e.g. [88] where it is assumed that every species could be identified with a fixed probability (for sampling of higher taxa only, see [89]). The second issue, which is more serious, concerns the nature of the species-level birth-death model we have been using. For the past, this model is an appropriate representation of the outcomes of the evolutionary process, and abundance of species is an appropriate measure of diversity. However, as we approach the present, the validity of this model arguably breaks down, as species lose their singular identity and become more accurately represented by individuals, sub-populations and nascent new species (see e.g. [90] [16]). This break-down is at least partly likely to account for the apparent lack of an observed POTPr in molecular phylogenies, which has been attributed to various types of background rate heterogeneity [16]. Although this topic is clearly an active area of research, one approach as far as detection of the LCE is concerned would be to compare the relative *present* sizes of otherwise equivalent clades of similar ages. The LCE would predict that their early rates of lineage diversification and the size of their subsequent slowdown would be proportional to their current clade size. The general stochasticity of the whole process can of course lead to a very wide range of possible outcomes and a suitably large sample of clades would be required in order to reliably detect the effect. Simulation of large numbers of clades, with a range of both POTPa and LCE, and taking into account the possible disturbing effects of mass extinctions, may assist in fully understanding the range of possible outcomes of rate heterogeneity that can arise from homogeneous models.

The intersection of background rate heterogeneity and survivorship biases raises important issues about the generalisability of theories about diversification. Clades that survive until the present day are biased by the POTPa, and of those, the large clades will be further biased by the LCE. Thus, large living clades represent a very unrepresentative sample of clades in general, and their features will not be universal to the entire population of clades that have been generated by the evolutionary process. These unusual clades can of course be modelled with specific models that describe the rate shifts that must exist in them. Such models could be used to systematically generate similar clades, but would likely fail to generalise to the more numerous, smaller and non-surviving clades. Thus, analysing only large, surviving clades to the exclusion of smaller and extinct clades will not demonstrate whether the properties of the studied clades results from survivorship biases, exogenous rate variation, or both. Indeed, the very simplicity of birth-death models and their powerful application suggests that the data we have are not in general sufficient to distinguish between the various ecological and evolutionary events that eventually give rise to them. As Nee remarked some years ago: “It is well known that completely different mechanisms can generate the same pattern: the distribution of parasitic worms among people is the same as the distribution of word usage in Shakespeare ― the negative binomial. This means that the patterns themselves cannot inform us about mechanism and some other techniques are needed” [91].

## 9 Summary

In this paper we have explored the patterns of diversification that can be generated by a retrospective view of a purely homogeneous process of diversification. These patterns can be substantial and highly non-homogeneous, and it is essential to understand these “null” hypotheses before considering causal explanations for any residuals (c.f. [14]). Patterns of diversification through time have been much discussed in the literature (e.g. [72]), with a common pattern being seen that diversification rates are high at the beginning of major evolutionary radiations, in both raw diversity counts and lineages through time plots. Various mechanisms for such effects have been proposed (such as filling empty ecological niches or unusual or flexible developmental evolution). The question that the analysis above poses is, however: are such patterns inevitably generated by the push of the past and/or the large clade effect? We have shown that the POTPa is strongest when background extinction rates are high, and that in likely scenarios for the evolution of large clades, it eventually accounts for nearly all of modern diversity. Furthermore, the POTPa impacts other many aspects of diversification dynamics, including recovery from mass extinctions. Indeed, the universality of such processes extends beyond evolutionary biology, with similar patterns being observed, for instance, in the size- or age-dependent growth of companies (see e.g. [92] and references therein). Even under homogeneous models, large clades can be generated at the edge of likely distributions that possess another characteristic, the “large clade effect”, which generates distinctive patterns in phenotypic and molecular evolution.

Harvey et al. (1994) [18], when briefly describing the POTPa, commented that “If these statistical effects are not fully appreciated, it could be tempting to misinterpret such a higher early slope as evidence for lineage birth rates being higher, and/or lineage death rates being lower, at earlier times” (p. 526). Here we have attempted to quantify both the size of, and controls on this effect, and to show just how important it in patterns of changes of rates of evolution through time including: dependency of rates of diversification on diversity; initial bursts of diversification at the origin of clades and the effects of mass extinctions. Although it seems natural to take the history and diversification of large and ultimately successful clades such as the arthropods as proxies for evolutionary radiations as a whole (e.g. [93] [67]) including after mass extinctions, our analysis shows this to be particularly fraught with difficulties: the history of life was written by the victors.

